# Identification of peptide candidate against COVID-19 through reverse vaccinology: An immunoinformatics approach

**DOI:** 10.1101/2020.07.01.150805

**Authors:** Rohit Pritam Das, Manaswini Jagadeb, Surya Narayan Rath

## Abstract

Novel corona virus disease 2019 (COVID-19) is emerging as a pandemic situation and declared as a global health emergency by WHO. Due to lack of specific medicine and vaccine, viral infection has gained a frightening rate and created a devastating state across the globe. Here the authors have attempted to design epitope based potential peptide as a vaccine candidate using immunoinformatics approach. As of evidence from literatures, SARS-CoV-2 Spike protein is a key protein to initiate the viral infection within a host cell thus used here as a reasonable vaccine target. We have predicted a 9-mer peptide as representative of both B-cell and T-cell epitopic region along with suitable properties such as antigenic and non-allergenic. To its support, strong molecular interaction of the predicted peptide was also observed with MHC molecules and Toll Like receptors. The present study may helpful to step forward in the development of vaccine candidates against COVID-19.

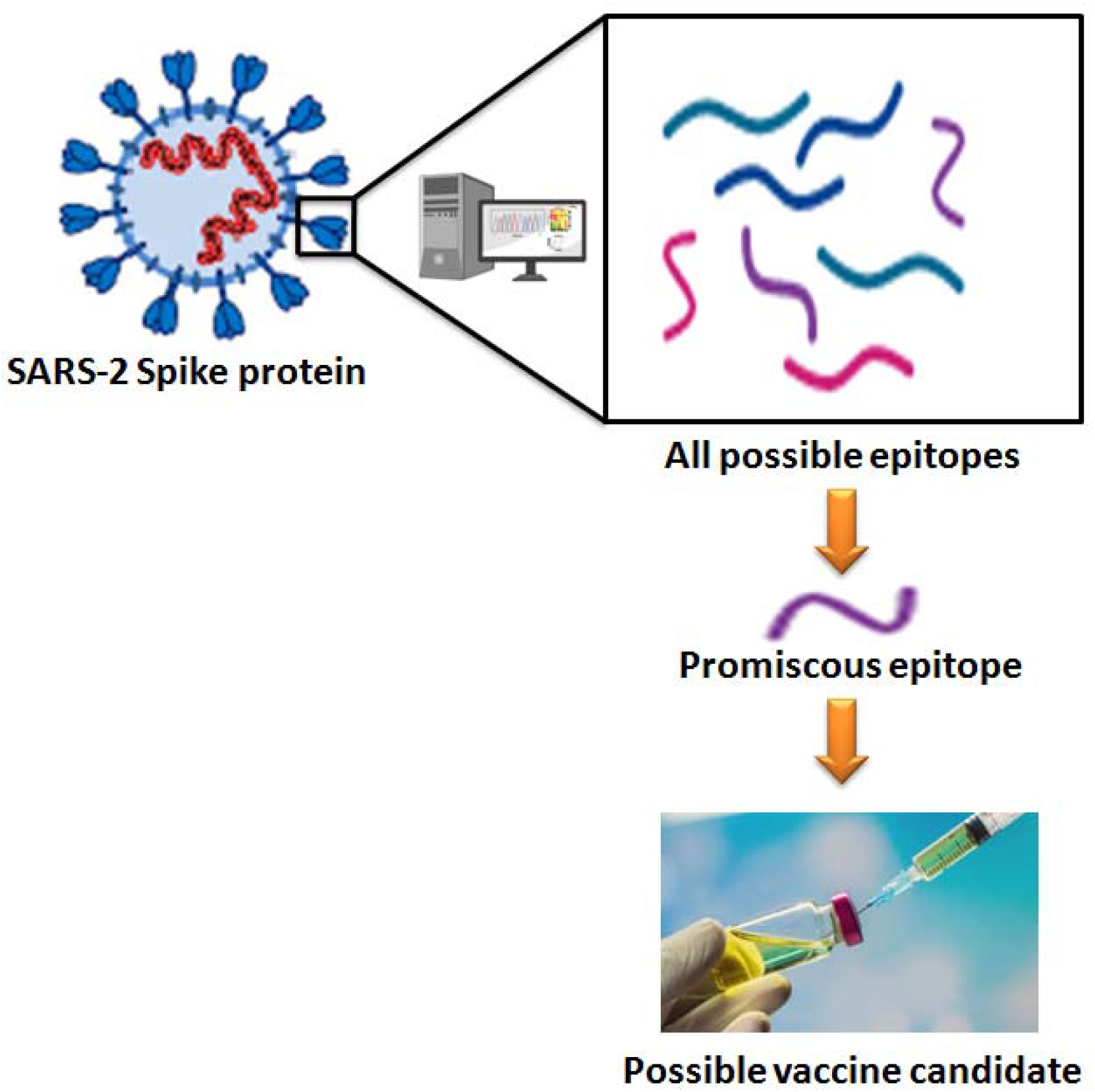

## Introduction

The disease COVID-19 outbreak caused due to emergence of novel severe acute respiratory syndrome corona virus 2 (SARS-CoV-2) [1]. According to WHO, the novel corona virus has affected more than 5.5 million people with a fatality of 3, 46,969 across the globe as of May 25, 2020. Respiratory droplets, direct contact and fecal-to-oral transmission are conventional routs for SARS-CoV-2 [2]. The symptoms of SARS-CoV-2 infection include fever, dry cough, shortness of breath, runny nose and sore throat [3, 4]. The rate of transmission and death is gaining severity due to ignorance of specific drug and vaccine against it.

SARS-CoV-2 is a positive-sense single-stranded RNA virus and its genome is around 29.7 kB long with twelve putative open reading frames (ORFs) that encode different structural and non-structural proteins. The first SARS-CoV-2 (Wuhan-Hu-1) genome was successfully sequenced and submitted to GenBank on January 5, 2020 (Accession no. MN908947.3) [3]. One-third of the genome is responsible for coding the structural proteins in SARS-CoV-2, namely, spike (S), envelope (E), membrane (M), and nucleocapsid (N) of SARS-CoV-2 are potential antigen for neutralizing antibody preparation and may be prospective therapeutics [5]. After entering into host body, the virion attaches to the host cell membrane and the viral Spike protein S1interacts with a functional host cell receptor known as angiotensin-converting enzyme 2 (ACE2). Thereafter, Spike protein S2 mediates the fusion of the virion and cellular membranes by acting as a class I viral fusion protein [6]. During this phase, the protein attains at least three conformational states: pre-fusion native state, pre-hairpin intermediate state, and post-fusion hairpin state [7]. As SARS-CoV-2 S glycoprotein is surface-attached and has potentiality to initiates the infection thus could be a promising vaccine target. In this connection, epitope based peptide design have remarkable privilege than conventional vaccine development. Peptide based vaccine are most popular since they are specific, generate long lasting immunity, able to avoid undesirable immune responses and are reasonably cheaper [8]. In addition, epitope based vaccine design has been aided by robust computational techniques [9]. Therefore, authors have focused on discovery of epitope from SARS-CoV-2 S glycoprotein.

The T-cell epitopes are typically peptide fragments, whereas the B-cell epitopes can be proteins, lipids, nucleic acids or carbohydrates [10, 11, 12]. Based on literature, the peptide is considered sufficient for activation of the appropriate cellular and humoral responses as it is the fragment of antigenic protein [13, 14]. Here we have identified peptide as vaccine candidate as the peptide vaccines are comparatively easy for production, chemically stable, and absence of infectious potential [15]. The present study would throw lights on vaccine development against COVID-19.

## Materials and method

### Sequence retrieval

Protein sequence of S glycoprotein was retrieved from UniProt (ID: P0DTC2) database [16]. The Gene name is “S” which belongs to Severe acute respiratory syndrome corona virus 2 (2019-nCoV) (SARS-CoV-2).

### B-cell epitope prediction

The B-cell epitopic regions present in SARS-CoV-2 S protein were identified using BcePred prediction server (https://webs.iiitd.edu.in/cgibin/bcepred/) [17]. It helped to predict linear epitopes from S protein sequence using physico-chemical properties.

### T-cell epitope prediction

MHC binding prediction includes the prediction of binding sites for both CD4+ and CD8+. The IEDB analysis resource (http://tools.iedb.org/main/) predicts specific T-cell epitopes to bind with MHC class I molecules along with IC50 (half maximal inhibitory concentration) values. Similarly, it employs different methods to predict MHC Class II epitopes, including a consensus approach which combines NN-align, SMM-align and combinatorial library methods [18].

### Interaction with MHC molecules

The crystal structure of HLA-B*35:01 (PDB ID: 4LNR) presenting MHC class I molecule in complex form with the peptide (RPQVPLRPMTY) was retrieved from PDB database [19]. Similarly, as representative of MHC-II, the crystal structure of HLA-DR1 (PDB ID: 1T5X) in complex form with a synthetic peptide (AAYSDQATPLLLSPR) was retrieved from PDB. PepFold server [20, 21] was used to build the tertiary structural model of predicted peptides. Molecular docking was performed between the predicted peptides and MHC representative structures using PatchDock web server [22, 23, 24].

### Antigenicity and allergenicity prediction

Determination of antigenic and allergenic properties are two important factor related to peptide based vaccine designing. The antigenicity of predicted peptides was calculated using VaxiJen tool [25] with the cut off value 0.4. AllerTOP v. 2.0 [26] and AllergenFP v.1.0 tool [27] was used to predict allergenic property of predicted peptides.

### Physico-chemical properties prediction

In order to find the molecular properties of predicted peptides, Innovagen’s peptide calculator was used. It makes calculations and estimations on physiochemical properties like peptide molecular weight, peptide extinction coefficient, peptide net charge at neutral pH and peptide iso-electric point.

### Interaction with Toll-Like Receptors (TLRs)

Crystal structures of human Toll-Like receptors such as TLR2 (PDB ID: 6NIG) and TLR4 (PDB ID: 4G8A) was extracted from PDB and subjected for structural preparation. Interaction of both TLR2 and TLR4 structures with the predicted peptides were performed using PatchDock web server [22, 23, 24].

## Results and Discussion

### *Sequence retrieval and antigenic property prediction of* SARS-CoV-2 ‘*S’ protein*

The complete sequence of SARS-CoV-2 S protein (UniProt ID: P0DTC2) is of 1,273 amino acids length. Average antigenic propensity was calculated as 1.0416 from 63 antigenic determinants within its primary sequence. The antigenic plot (**Figure 1**) between amino acid residues with respect to propensity strongly established S protein antigenicity.

**Figure 1:**
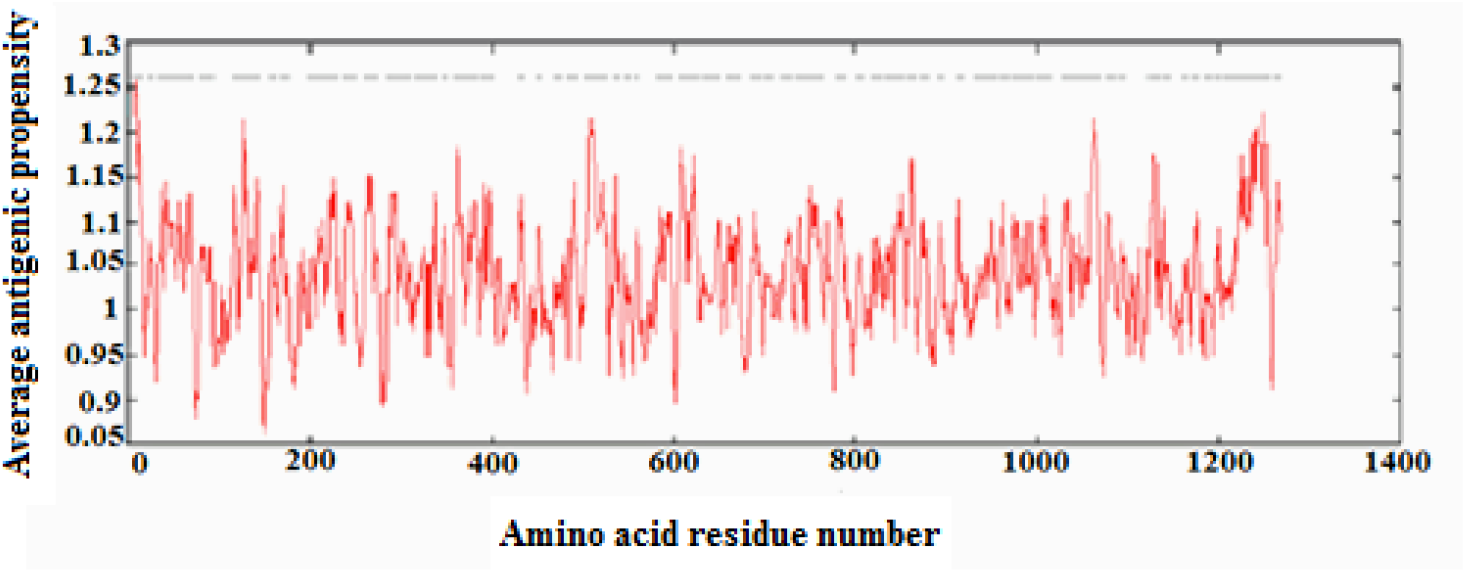
Average antigenic propensity plot of SARS-CoV-2 spike protein

### B-cell and T-cell epitope prediction

Prediction of B-cell epitope is a crucial step in epitope base vaccine design [28]. Linear B-cell epitope region of SARS-CoV-2 S protein was identified with the help of its physicchemical properties. Similarly the tabular representation of B-cell epitope reveals the amino acid at 84, 85, 91, 92 in pep-9 and 521, 522, 523, 524, 525 in pep-15 belongs to the B-cell epitopic region (**Figure 2**).

**Figure 2:**
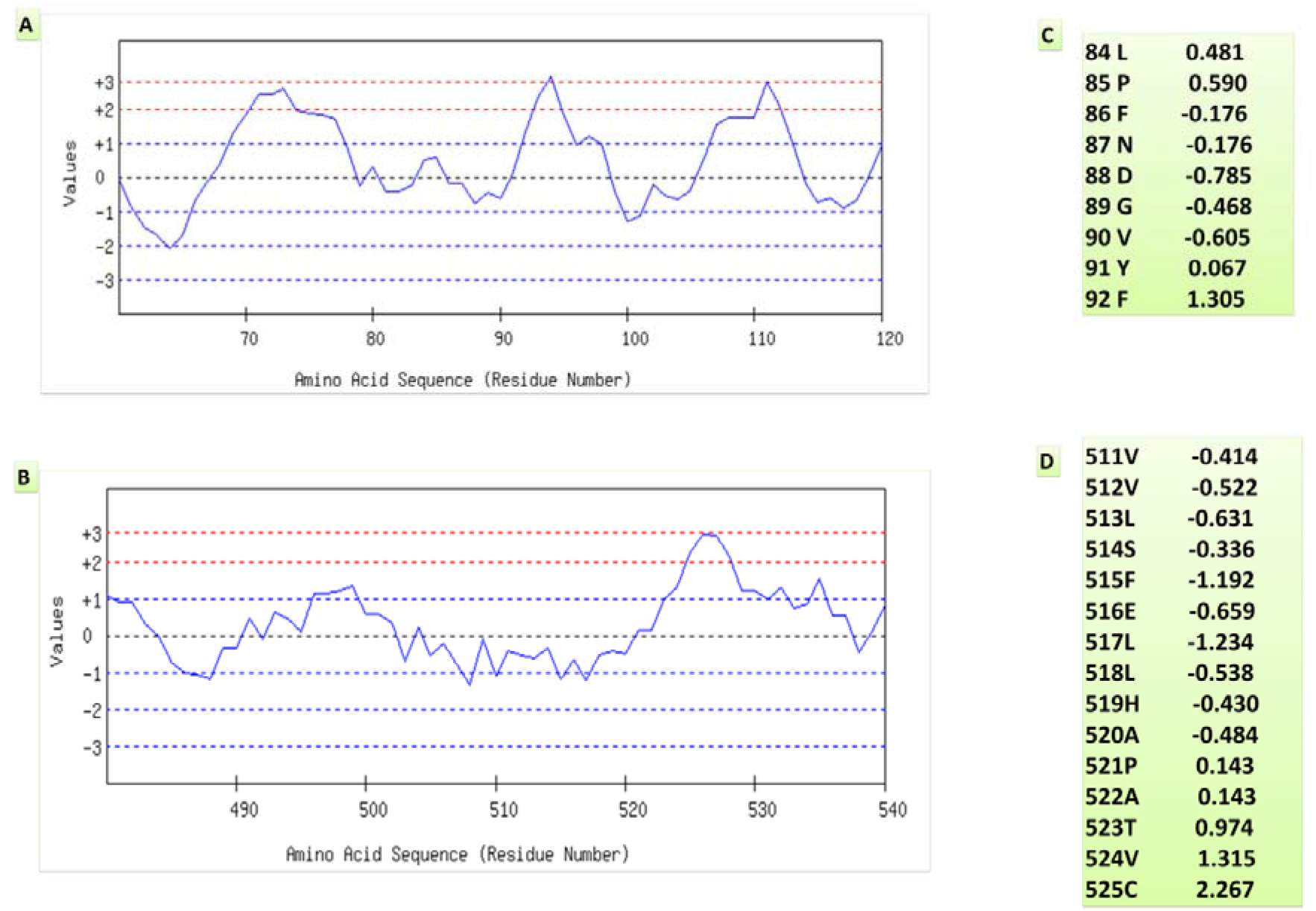
B cell epitope regions for predicted pep-9 and pep-15 for SARS-CoV-2 spike protein. (A) and (B) represents the graphical representation of selected amino acid regions. (C) and (D) represents the epitopic scoring of each amino acid from predicted peptides.

T-cell epitope prediction is also known as a well establish method in vaccine discovery. Here we identified more than 100 T-cell epitopes with different percentile rank for the whole protein length. Basically, peptides with least percentile rank indicate their higher binding affinity against T-cell. therefore only five peptides within 0.01 to 0.03 percentile rank were found suitable for CD8+. Among them, four peptides with amino acid position 39-47, 45-53, 39-47 and 27-35 were suitable for HLA-B27 molecule where as the only peptide (24-32) suitability attached with HLA-B35 molecules. Further, the 9-mer peptide (Pep-9) LPFNDGVYF (84-92 amino acid) was represented for both B-cell and CD8+ T-cell therefore, selected for this study. Similarly, in case of CD4+ T-cell epitope prediction, out of five top ranked peptides (0.01 percentile) none of them were represented by B-cell thus discarded. Furthermore, one 15-mer (511-525 amino acid) peptide (VVLSFELLHAPATVC) (Pep-15) was identified suitable for both B-cell and CD4+ T-cell epitope (percentile rank 0.03) region.

### Molecular interaction between peptides and MHC molecules

Structure of MHC-I molecule namely HLA-B*35:01 (PDB ID: 4LNR) was identified as most suitable for Pep-9 (LPFNDGVYF). Similarly, MHC-II molecule HLA-DR1 (PDB ID: 1T5X) was found suitable for Pep-15 (AAYSDQATPLLLSPR). The attached synthetic peptides were removed from their respective crystal structures in order to free the peptide binding sites. Molecular docking was performed between representative structures of MHC molecules and suitable peptide candidates in order to inspect their molecular interaction. Independent dock run was performed in both of the system which produced ten binding complex for each of docking. Five top ranked docked complexes were reported with atomic contact energy (ACE) scores (**Table 1**). The ACE plays an important role in peptide binding efficacy which is known to be negative in strong interaction. Therefore, model 5 i.e. HLA-B*35:01-Pep9 (dock score: 7394; interaction area: 940.60Å; ACE: −191.22 kcal/mol) and model1 i.e. HLA-DR1-Pep15 (dock score: 8852; interaction area: 1152.50 Å; ACE: −236.54kcal/mol) were selected for Pep-9 and Pep-15 peptide binding for further analysis.

**Table 1:**
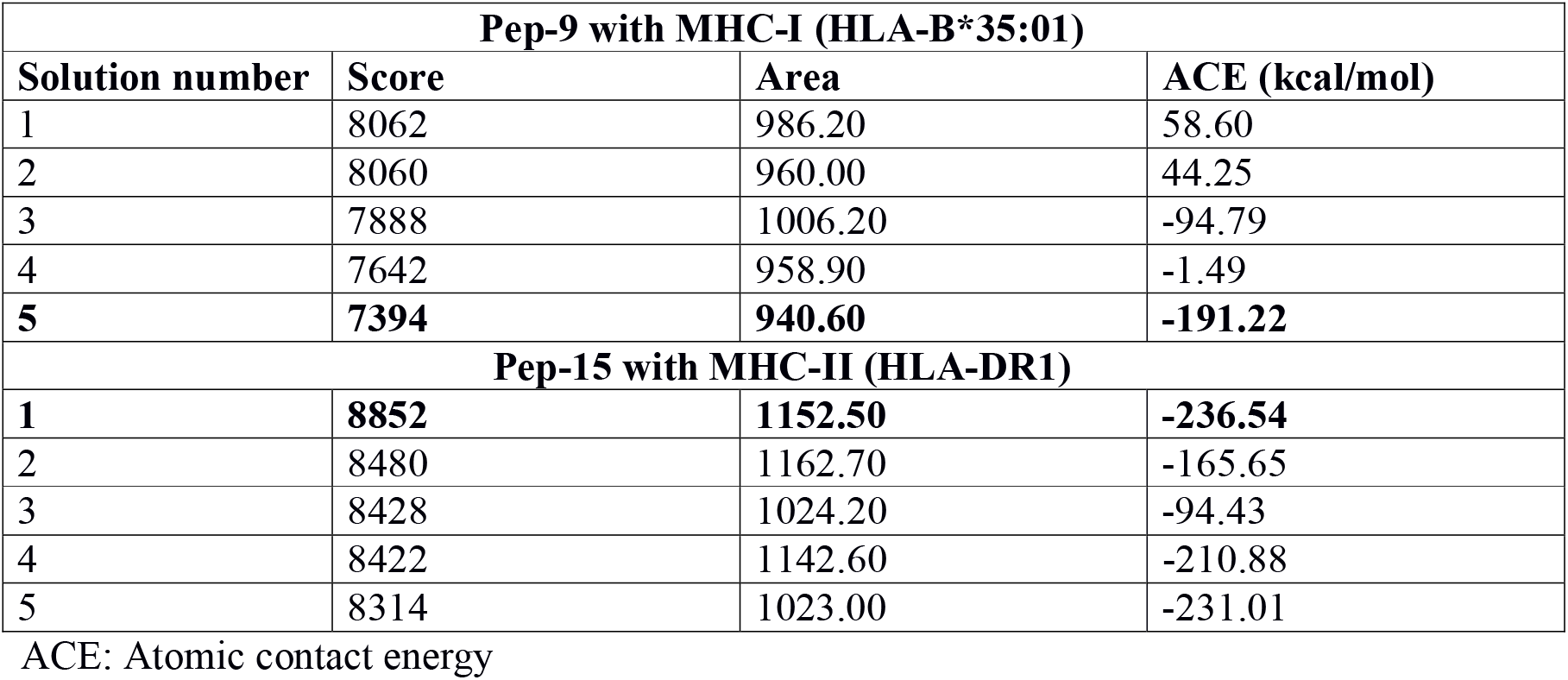
Molecular docking scores between predicted peptides and MHC molecules were reported.

Interestingly, Pep-9 was interacted within the known peptide binding site (LYS58, GLU212, ALA211, SER4, ASP29 and ASP30) of the HLA-B*35:01 structure (PDB ID: 4LNR) where as Pep-15 was bound within a different functional site (THR77, ARG71, GLU55, GLN57 and ASN62) of HLA-DR1 structure (PDB ID: 1T5X) (**Table 2, Figure 3, Figure 4**) which signified about the suitability of Pep-9 as a potential vaccine candidate.

**Table 2:** Polar contacts between predicted peptides and MHC molecules were reported.

| Pep-9 and MHC-I (HLA-B*35:01) |  |
| --- | --- |
| Interacting amino acids | h-bond distance (Å) |
| LYS 58 | 4.5 |
| SER 4 | 3.4 |
| ASP 29 | 2.7 |
| ASP 30 | 2.5 |
| ALA 211 | 2.5 |
| GLU 212 | 4.3 |

| Pep-15 and MHC-II (HLA-DR1) |  |
| --- | --- |
| GLU 55 | 1.6 |
| GLN 57 | 3.4 |
| ASN 62 | 3.2 |
| ARG 71 | 3.4 |
| THR 77 | 3.0 |

**Figure 3:**
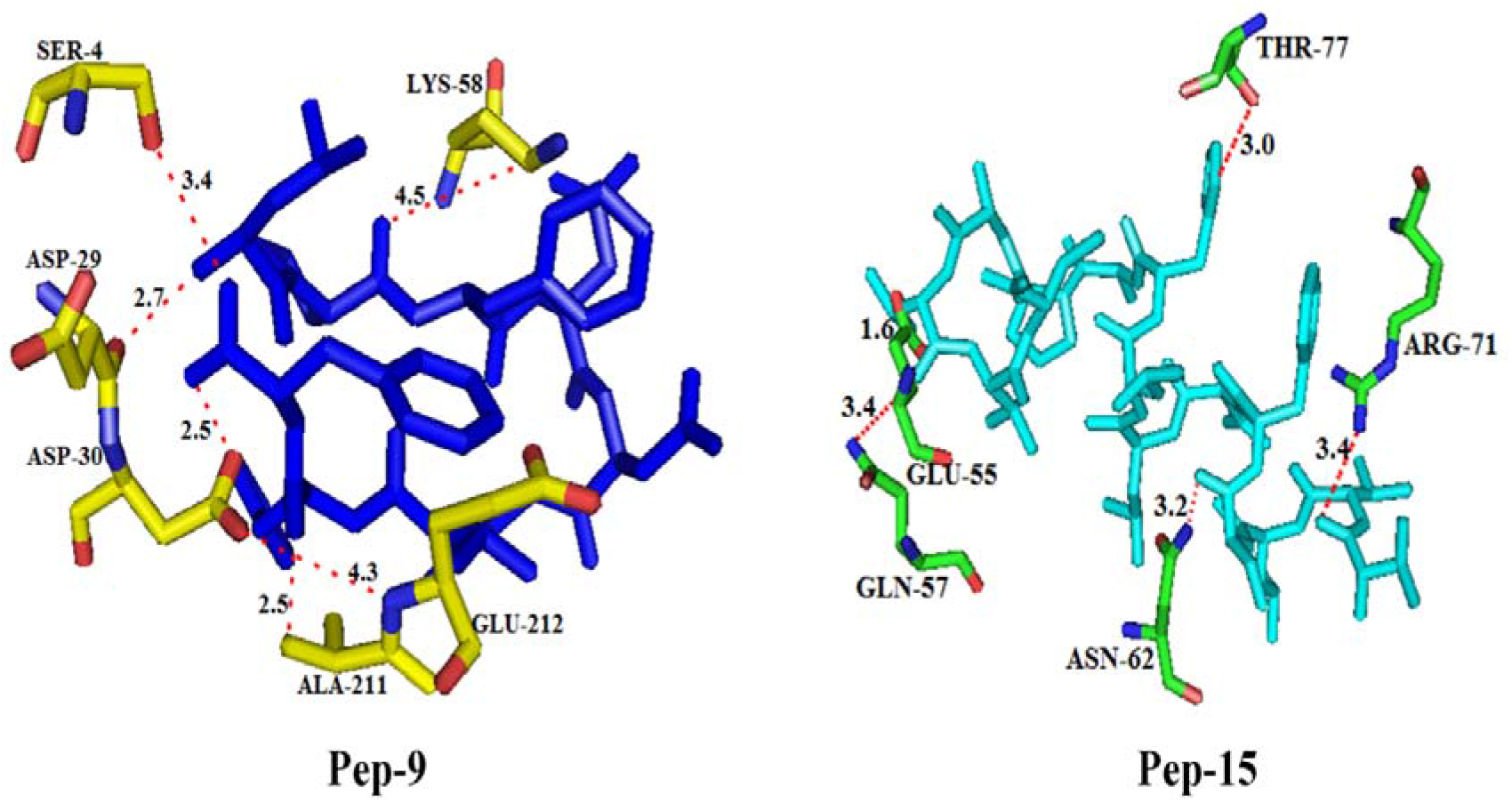
Atomic interactions between predicted peptides with MHC molecules were deciphered.

**Figure 4:**
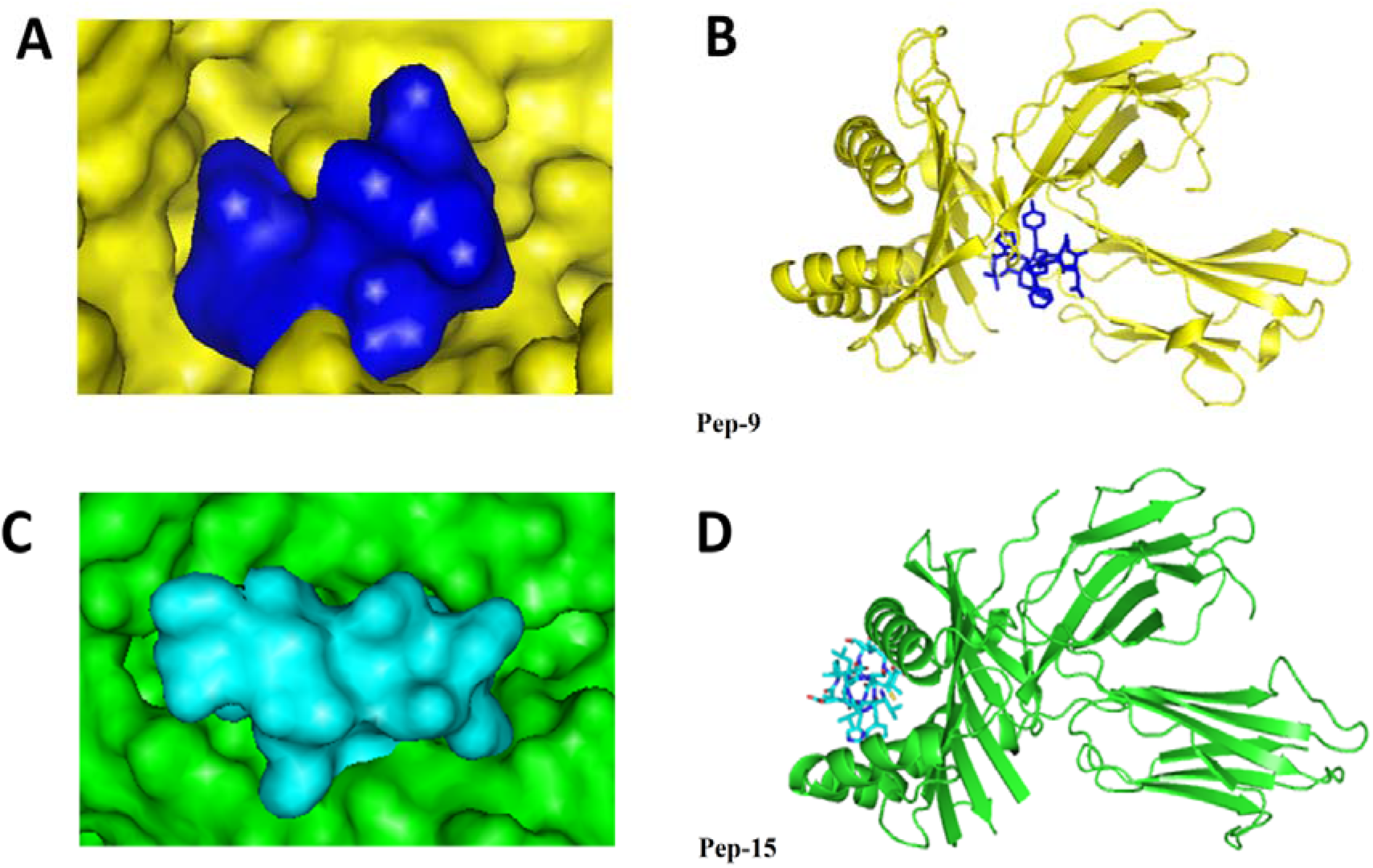
Docked peptides were visualized within functional sites of MHC molecules. (A, B): Binding pattern of Pep-9 (blue color) and MHC-I (HLA-B*35:01) (yellow color); (C, D) Binding pattern of Pep-15 (cyan color) and MHC-II (HLA-DR1) (green color)

### Analysis of antigenic, allergenic and physico-chemical properties of predicted peptide candidates

Antigenic peptides increase the immunogenicity of vaccine candidate therefore prediction of antigenic property is an essential step in immunoinformatics based vaccine design. In this study, both of the peptide Pep-9 (antigenic score: 0.5593) and Pep-15 (antigenic score: 0.8618) was showed above threshold value (0.4) thus established as suitable antigenic peptides. In addition, both of the peptides were predicted as non-allergen by AllerTOP v. 2.0 and AllergenFP v.1.0 tools. Further, molecular weight of Pep-9 and Pep-15 were calculated as 1071.18 g/mol and 1598.91 g/mol respectively. To its support, hydropathy calculation determined about the hydrophobic nature of both peptides (**Figure 5**). However, overall analysis confirmed about the effectiveness of both Pep-9 and Pep-15.

**Figure 5:**
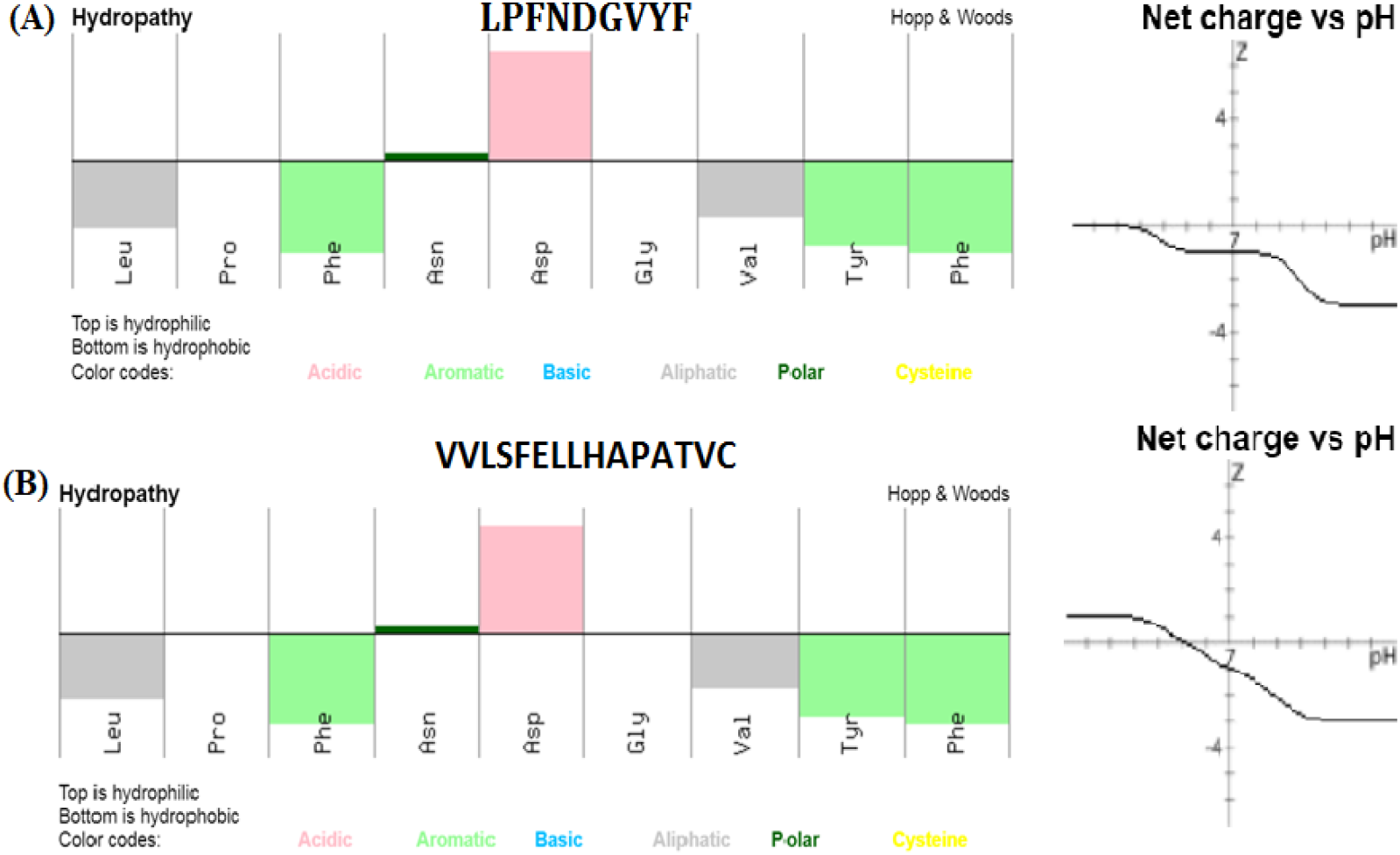
Physico-chemical properties of selected peptides. (A) Pep-9 (B) Pep-15.

### Interaction of peptides with TLR2 and TLR4

The potential role of Toll-Like receptor mediated host-parasites interaction is well established. Particularly, TLR2 and TLR4 are well known mediums for interaction between filarial parasites and host innate immune system [11, 14, 15]. Therefore, in this study we have studied the interaction between predicted peptides (Pep-9 and Pep-15) and Toll-Like receptors (TLR2 and TLR4). The result suggested Pep-9 more strongly interacted with TLR2 than TLR4 (**Table 3**, **Figure 6**). Further, strongest binding affinity (ACE score) was observed between TLR2-Pep15 docked complex (**Table 3**, **Figure 6**). Moreover, Pep-9 is found more significantly binding with both TLR2 and TLR4 (**Table 3**, **Figure 6**).

**Table 3:** Molecular interaction scores between predicted peptides with Toll-Like Receptors

| Interaction | Score | Area | ACE |
| --- | --- | --- | --- |
| **TLR2-Pep9** | **7288** | **971.10** | **-305.36** |
| TLR4-Pep9 | 5950 | 680.20 | -28.51 |
| **TLR2-Pep15** | **8306** | **1031.00** | **-451.70** |
| TLR4-Pep15 | 1075 | 11.53 | 11.53 |

ACE: Atomic contact energy

## Conclusion

This study is focused on the prediction of effective epitopes from Spike protein of SARS-CoV-2. Among all possible epitopes, Pep-9 and Pep-15 were claimed as more effective candidates. Again, it was observed that the Pep-9 as more suitable than Pep-15 in order to be a possible vaccine candidates. Further, *in vitro* and *in vivo* validation is required to confirm the prediction. Overall, this study would be informative towards new vaccine development for prevention of widespread COVID-19.

## Acknowledgment

We are thankful to Dr. Pawan Kumar Agrawal, Vice chancellor, Odisha University of Agriculture and Technology, Bhubaneswar for his moral support and valuable suggestion.

## Conflict of interest

The authors declare no competing interest.

## Notes

### Competing Interest Statement

The authors have declared no competing interest.

